# A Dynamic Efficient Sensory Encoding Approach to Adaptive Tuning in Neural Models of Visual Motion Processing

**DOI:** 10.1101/2021.06.17.448823

**Authors:** Scott T. Steinmetz, Oliver W. Layton, Nate V. Powell, Brett R. Fajen

**Affiliations:** Cognitive Science Department, Rensselaer Polytechnic Institute; Computer Science Department, Colby College

**Keywords:** efficient coding, adaptation, neural modeling, optic flow

## Abstract

This paper introduces a self-tuning mechanism for capturing rapid adaptation to changing visual stimuli by a population of neurons. Building upon the principles of efficient sensory encoding, we show how neural tuning curve parameters can be continually updated to optimally encode a time-varying distribution of recently detected stimulus values. We implemented this mechanism in a neural model that produces human-like estimates of self-motion direction (i.e., heading) based on optic flow. The parameters of speed-sensitive units were dynamically tuned in accordance with efficient sensory encoding such that the network remained sensitive as the distribution of optic flow speeds varied. In two simulation experiments, we found that model performance with dynamic tuning yielded more accurate, shorter latency heading estimates compared to the model with static tuning. We conclude that dynamic efficient sensory encoding offers a plausible approach for capturing adaptation to varying visual environments in biological visual systems and neural models alike.

## INTRODUCTION

Biological visual systems are remarkably adaptive to a wide range of visual environments and rapidly varying conditions, enabling animals to maintain perceptual contact with their surroundings regardless of whether they are familiar or novel, static or fluctuating. Although biological agents often encounter dramatic shifts in stimulus values as they move from one context to another, they rarely experience meaningful degradation in perception or performance. Understanding the principles and mechanisms that allow for such robustness is critical for both advancing our theoretical understanding of the visual system and informing the development of computational neural models of biological vision. The aim of this study is to explore and evaluate the idea that adaptation can be modeled as continuous changes in the mapping of neuronal inputs to outputs. We tie these changes in response to changes in the distribution of a stimulus variable in accordance with the principles of efficient sensory encoding (Ganguli & Simoncelli, 2014), extending an approach that was originally envisioned as a “set-it-and-forget-it” method for parameter tuning to scenarios that demand rapid adaptation in real-time.

Visual adaptation encompasses a broad set of phenomena that extend across multiple timescales (Wainwright, 1999): the visual system is broadly shaped by its environment via evolutionary processes (Attneave, 1954; Barlow, 1961), it is attuned to local natural scene statistics through experience (Simoncelli, 2003; Simoncelli & Olshausen, 2001), and it rapidly adapts to current stimuli (Webster, 2015; Werblin, 1974). Adaptation has been studied extensively at both the perceptual and neural levels. The perceptual consequences of such adaptation include aftereffects, which are perceptual vestiges of adaptation to recently presented stimuli, as well as decreases in discrimination thresholds for recently experienced stimuli. At the neural level, adaptation is reflected in shifts in the firing rates of cells in response to prolonged exposure to a constant stimulus. Such effects have been observed across a variety of stimulus properties, including motion (Addams, 1834; Bartlett et al., 2018), light intensity, and orientation (Carandini et al., 1998; Dragoi et al., 2000; Gutnisky & Dragoi, 2008; Müller et al., 1999), and in both human and non-human animals (Howard et al., 1987; Laughlin, 1989; Shapley & Enroth-Cugell, 1984).

Adaptation in the visual system can be understood as a consequence of optimal information transmission, which captures how neural connections adjust to more efficiently encode relevant sensory variables (Barlow, 1990; Brenner et al., 2000; Laughlin, 1989; Wainwright, 1999). One highly influential theory that ties these concepts together is the efficient coding hypothesis (Attneave, 1954; Barlow, 1961), which is rooted in the idea that the information available to early sensory neurons is highly redundant and that such neurons distill from the flood of information the most relevant perceptual properties (Olshausen & Field, 1997). As the agent moves and its surroundings change, the statistics of naturally occurring stimuli undergo variations, which in turn require neural systems to adapt in order to maintain efficiency. Indeed, a static mapping from input to output is necessarily less efficient at extracting and encoding a signal than a dynamic mapping that adapts to new conditions.

Such adaptation to changing stimuli has long been predicted to have perceptual consequences, as sensitivity to more relevant information increases at the cost of lower sensitivity to less relevant information (Laughlin, 1989). Neurophysiological studies have found that post-adaption changes in neural activity that reflect coding efficiency are correlated with perceptual adaptation, for example in macaques viewing variously oriented gratings (Gutnisky & Dragoi, 2008). This has also been borne out in behavioral studies where humans have exhibited improved discrimination to recently encountered optic flow and degraded discrimination to older flow patterns (Durgin & Gigone, 2007). It is in this sense that adaptation can be understood as a consequence of coding efficiency.

### The present study

The goal of this study is to explore a principled self-tuning mechanism for neural models of early visual areas that captures how biological systems adapt to wide ranges of conditions while maintaining precision and accuracy. Our solution extends efficient sensory encoding (Ganguli & Simoncelli, 2014) which is itself built upon the efficient coding hypothesis (Attneave, 1954; Barlow, 1961). Efficient sensory encoding offers a principled approach to defining the parameters of tuning curves for N neurons such that they optimally encode a given distribution of stimulus values. In Ganguli and Simoncelli’s formulation, this distribution was based on the values of stimulus variables found in the agent’s environment. Their approach provides a mathematically precise means of codifying and optimizing tuning curve selection such that the neural population produces activity and discrimination thresholds similar to those seen in previous studies. However, it also assumes a static distribution of stimulus variables, which yields a static neural tuning.

We introduce dynamic efficient sensory encoding (DESE) as a mechanism for adaptive neural tuning to rapidly shifting distributions of stimulus variables. DESE extends the approach to tuning curve selection introduced by Ganguli and Simoncelli (2014) to define attunement based on recently detected sensory information, dynamically adapting with changes in the distribution of stimulus values. This provides a mathematical basis for continual and automated parameter tuning for early visual neurons that optimizes the encoding of target variables given the stimuli the agent recently encountered, instead of the statistical average for the environment. Beyond being a potential biological mechanism employed by early sensory neurons to capture adaptation effects such as those observed in the H1 neuron in flies (Brenner et al., 2000), this approach could benefit modelers who build computational neural models of vision and aspire to efficiently simulate the behavior of a large number of units with finite computing resources. Dynamic attunement also has the advantage of allowing for adaptation to the current sensory environment without modeler intervention, which is important both for avoiding overfitting and for practical applications where manual parameter selection is both unprincipled and time consuming.

We explore this solution in the context of an existing neural model, which provides us with a concrete example and allows for performance comparison of DESE against previous static approaches. Specifically, we discuss the dynamic tuning of speed-sensitive cells in the middle temporal (MT) area of the competitive dynamics (CD) model of optic flow processing in the primate dorsal stream (Layton & Fajen, 2016b, 2016a). In the next few sections, we briefly introduce the reader to the topic of heading perception, summarize the CD model, and explain how we implemented dynamic efficient sensory encoding within the CD model. We then report the results of a set of simulation experiments that explore the improvement in heading estimation when dynamically attuning to the stimulus distribution.

### Optic Flow and the Perception of Heading

Humans routinely navigate complex and dynamic environments without colliding with the other inhabitants and structures. By most accounts, the ability to successfully avoid obstacles and reach goals relies on sensitivity to one’s direction of self-motion or heading (Li & Cheng, 2011). Although multiple sensory modalities contribute to the perception of self-motion (Cullen, 2019; Greenlee et al., 2016), the ability to perceive where one is headed relative to potential goals and obstacles is predominantly driven by vision. In particular, heading perception is based on the structured patterns of optical motion that are induced by self-motion and known as optic flow (Browning & Raudies, 2015; Gibson, 1950; Warren et al., 2001). When the agent is traveling along a linear path through a static environment, the global optical motion radiates from a single point called the focus of expansion (FoE) that specifies the direction of self-motion. Indeed, humans can perceive their heading direction from optic flow with an accuracy of less than 1° of visual angle (Warren et al., 1988). Neurons that respond to global optic flow patterns and exhibit heading tuning have been found in various regions of the primate visual system, including MSTd (Duffy & Wurtz, 1991; Gu et al., 2006) and VIP (Britten, 2008; Chen et al., 2011).

### Competitive dynamics model

Research on the neural mechanisms involved in the processing of optic flow in the primate visual system has informed the development of computational neural models of heading perception (Browning et al., 2009; Beintema & Van den Berg, 1998; Lappe & Rauschecker, 1994; Perrone, 2012; Royden, 2002), including the competitive dynamics (CD) model (Layton & Fajen, 2016) used in the present study. The CD model comprises units that are organized in layers (e.g., V1, MT, MST) with response properties similar to those found in the corresponding areas of the primate brain (see Figure 1). Estimates of heading direction generated by the model are consistent with those of humans across a variety of scenarios, including in the presence of locally and globally discrepant optic flow resulting from factors such as independently moving objects and blowing snow, respectively (Layton & Fajen, 2016a, 2016c, 2017; Steinmetz et al., 2019). Such robustness is largely due to mechanisms such as spatial pooling and recurrent feedback that allow the model to make use of the temporally evolving global flow field rather than relying on an instantaneous snapshot of optic flow as prior models did (Layton & Fajen, 2016b).

**Figure 1:**
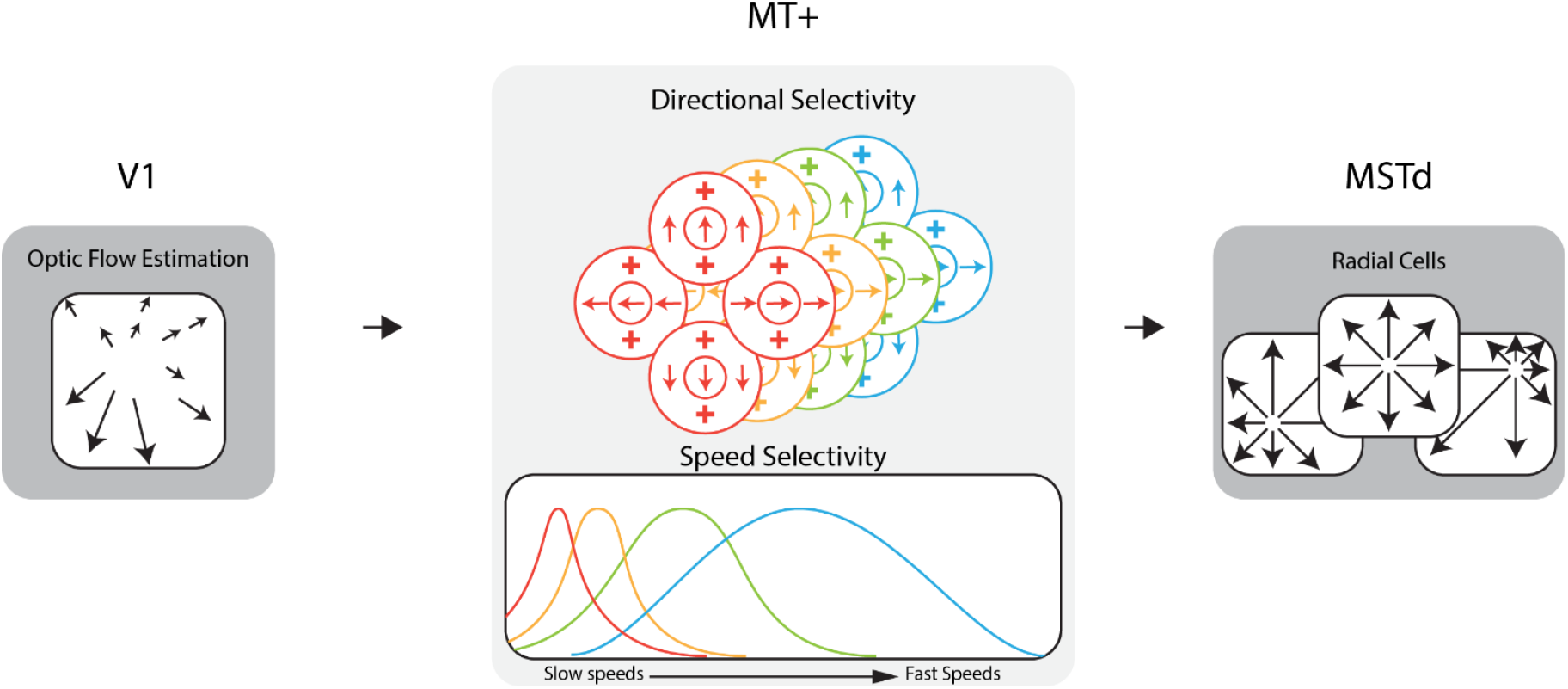
Diagram of the dorsal stream of the competitive dynamics model. The model takes optic flow as input for an approximation of the V1 feedforward signal. This optic flow drives area MT+ activity which encodes the optic flow signal within the population at each receptive field location (pixel). Model MT+ cells exhibit joint direction and speed tuning and pool motion inputs within the receptive field. In this figure, MT+ cells of a given color in the direction selectivity diagram have the speed tuning curve of that same color shown in the speed selectivity diagram. MT+ activity drives the activity of MSTd cells, which are sensitive to various patterns of flow, e.g. radial, laminar, etc.

### Speed selectivity in the CD model

In the CD model, as in the primate visual system, the local optical motion in any small region of the visual field is encoded in terms of the activity of MT neurons. Each individual unit exhibits tuning to a range of speeds and directions of motion. The model population consists of neurons tuned to all combinations of 24 directions and 7 speeds. For the purposes of the present study, speed is the more relevant of the two stimulus dimensions, so we will largely ignore variations in directional sensitivity from this point forward for the purposes of clarity. We will refer to a set of MT units with the same speed preference as a “layer”.

Each MT cell has a Gaussian-shaped tuning curve with two parameters: the mean (μ) which corresponds to the optic flow speed that maximally excites the unit, and the standard deviation (σ) which defines the range of speeds to which the unit responds. As such, the response of any given MT cell depends on how closely the optic flow speed matches the cell’s preferred speed, dropping to zero for speeds above and below the cell’s range of sensitivity.

The optical speed of an object or surface in the world is a function of the relative distance, direction, and motion between the agent and object (Longuet-Higgins & Prazdny, 1980). As such, during naturalistic self-motion, changes in the motion of the agent and the structure of the environment cause the distribution of optic flow speeds that one encounters to vary across a wide range. Although no single MT cell in the CD model is sensitive to the entire range of optic flow speeds, variations in the neuronal sensitivities allow for the population as a whole to be responsive to a wider range. Ideally, the number of unique speeds (or layers) to which MT neurons are tuned would be sufficiently large to fully sample with fine-grained precision the entire distribution of optic flow speeds that will be encountered in the world. Simply adding more units, however, introduces problems for both biological vision systems and neural models, which must be efficient in the use of resources. In the CD model, for example, MT units are tuned to common speeds to ensure uniform sensitivity across the visual field. As such, adding a unique speed tuning entails adding an entire layer of MT cells (65,536 units), which dramatically increases the computational cost of simulation. In the present study, area MT had seven different speed layers; that is, each MT unit could take on one of seven unique parameter value pairings (μ_i_, σ_i_, i = 1,…, 7).

The model’s overall sensitivity to motion depends not only on the number of layers but also on how well speed tuning curve parameters match to the input distribution of optic flow speeds. Because each layer is only sensitive to motion within a limited range (see <u>Movie 1</u>), selecting model tuning curve parameters that do not capture the range of speeds present in the optic flow input would result in regions of poor or absent sensitivity. This is a critical point because a poorly tuned model will haphazardly use only a portion of the available optic flow, potentially leading to sub-optimal performance. Conversely, when the tuning curves are optimally distributed along the stimulus space, the neurons encode the signal as efficiently as possible.

One might reason that tuning curves should uniformly tile the range of stimuli seen, as depicted in Figure 2a. However, if speeds do not arise with uniform probability in the optic flow (Figure 2b), a uniform tiling would result in equal precision between frequently observed stimuli and infrequently observed stimuli. A more efficient use of resources would be to recruit more neurons for signaling common stimulus values so that such stimuli can be encoded more precisely, and fewer neurons to signaling less common stimulus values (Figure 2d).

**Figure 2:**
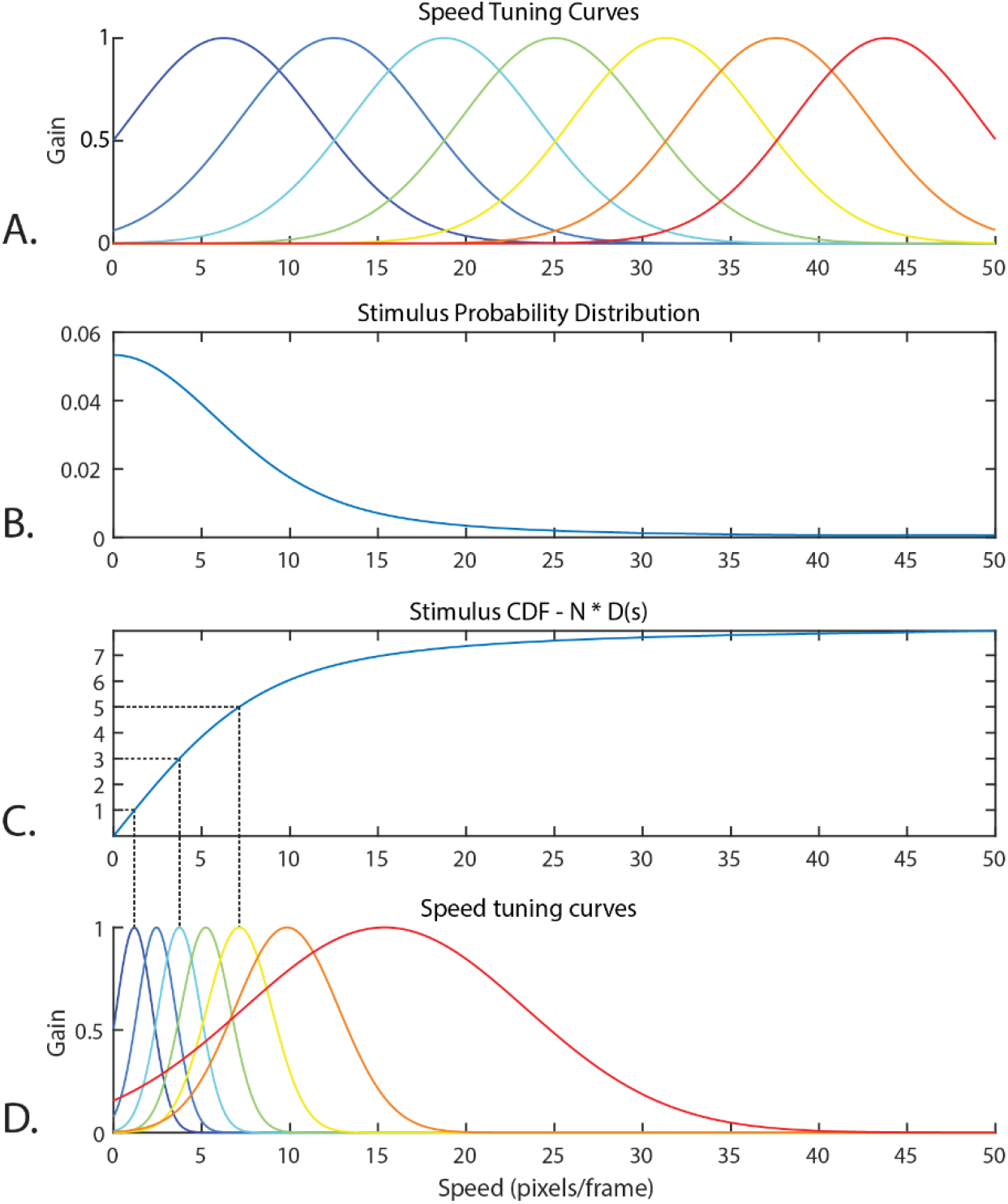
(A) Speed tuning curves that uniformly tile the stimulus range 0 to 50. Although the uniform tiling strategy could seem appealing, the distribution of optic flow speeds present in the input optic flow within the video dataset used in Experiments 1 and 2 of the present study (B) is clearly non-uniform. The uniform tiling strategy leaves much of the encoding capacity of the uniform tuning wasted on infrequently observed speeds. The tuning curves derived via efficient sensory encoding (D) allocate neural sensitivities based on the frequency that each speed arises in the stimulus, calculated via the (C) CDF of the probability distribution multiplied by the number of encoding neurons, N. In this way, encoding capacity (and computational cost) is optimally allocated for encoding this distribution.

### Efficient Sensory Encoding

Efficient sensory encoding (Ganguli & Simoncelli, 2014) offers a principled approach rooted in the efficient coding hypothesis for calculating the peak locations and widths of the optimal tuning curve for a given stimulus. More precisely, sensory variables are encoded to optimize the Fischer information across the neural population given a stimulus probability distribution. The derived tuning curves proportionately sample the stimulus probability distribution, resulting in narrow and densely packed tuning curves where the probability of that stimulus is high (e.g. where instances of that optic flow speed are very common) and broad, sparse tuning curves where the probability of that stimulus is low. This results in each speed cell capturing an approximately equal probability mass of the sensory distribution.

The following procedure describes how the distribution of the stimulus values in the environment was used to calculate the parameters (*μ* and *σ*) of the Gaussian tuning curves. The density function, d(s), for a given stimulus magnitude, s, with N layers in the speed cell population is defined as

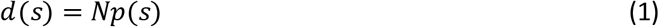

where p(s) is the probability distribution of the stimuli (see Figure 2b for an example). Note that multiplying by N causes the integral of d(s) to equal N. The peak location of the n^th^ speed cell’s tuning curve (*μ_n_*) is determined by evaluating where the cumulative distribution function (D) of the density function equals n:

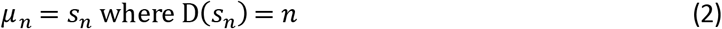

This process is illustrated in Figures 2c and 2d for N = 7 (the number of speed layers in the model used in the present study). The full width at half maximum (FWHM) of the n^th^ tuning curve is equal to the inverse of the density function evaluated at *s_n_*:

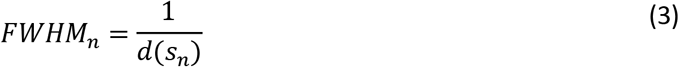

The FWHM is then used to calculate *σ_n_* as follows (for gaussian tuning curves):

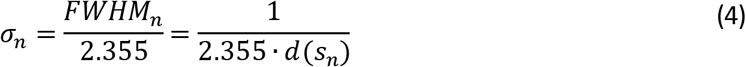

For an in-depth derivation of these equations, see Ganguli & Simoncelli (2014).

### Dynamic Efficient Sensory Encoding of Optic Flow

Efficient sensory encoding defines the ideal attunement for a given stimulus probability distribution but implicitly assumes that this attunement is stationary, which is not ideal for dynamic environments. The statistics of optic flow speeds vary not only across the visual field but also over time as the agent moves relative to objects and surfaces in the world (Calow & Lappe, 2007). Self-motion above a flat ground surface produces a smooth gradient of speeds proportional to the relative depth. By contrast, self-motion toward a flat wall produces in narrow range of optic flow speeds. Movement through realistic environments often produces a combination of constant speeds and gradients in the optic flow field. This is depicted in <u>Movie 2</u>, which shows the optic flow generated by self-motion through a simulated environment (Movie 2A) alongside a histogram that shows the distribution of optic flow speeds pooled across the entire image (Movie 2B). As the agent moves, changes in the direction and relative depth of surfaces as well as in self-motion speed and direction result in dramatic variations in the shape of the distribution over time.

In this study, we explore how the attunement process described in the previous section can be implemented on extremely rapid timescales (< 1 second). The model records and stores the distribution of optic flow magnitudes across the entire visual field over the past n frames. This defines the “ rolling time horizon” of the dynamic attunement, within which all flow is considered equally. The stored distribution of optic flow is then fed into the efficient sensory encoding equations defined in the previous section, and the speed cell tuning curves are adjusted according to the resulting values (Movie 2C).

## Methods

We conducted two simulation experiments to systematically test the potential benefits of dynamic efficient sensory encoding and to determine the effects of different time horizons. In Experiment 1, we compared dynamic tuning based on the evolving distribution of optic flow speeds observed over the last 10 frames to two static tunings, which served as control conditions. The first static tuning was based on a uniform distribution of optic flow speeds with a range of 0 to 30 pixels per frame. This range was chosen to span the most commonly scene optic flow reasonably (i.e. the body of the aggregate probability distribution which contained 96.37% of the observed optic flow). The second static tuning was based on the distribution of optic flow speeds aggregated across all frames of the entire test set of videos (described in the next paragraph). We refer to these three conditions as the Dynamic, Static – Uniform, and Static – Aggregate conditions, respectively. Experiment 2 investigated how performance with dynamic tuning depends on the number of previous frames across which flow vector magnitudes were aggregated to generate the optic flow distribution (i.e., the time horizon). Specifically, we compared the time horizon used in Experiment 1 (10 frames) with shorter (1 frame) and longer time horizons (30 frames).

We tested each tuning method on a set of 60 videos depicting self-motion through a high-fidelity virtual environment. Examples of the videos can be seen in <u>Movie 3</u>. The videos were generated using Microsoft AirSim in the Unreal game engine. We selected two distinct environments to provide a broad sample of natural and human-made scene structures: an outdoor residential neighborhood and the inside of a warehouse. The former included houses, parked cars, trees, and telephone poles with powerlines situated in residential zones and parks, various ground-surface textures (e.g., pavement, concrete, grass), and a blue sky with distant clouds. Most vantage points exhibited naturalistic depth variation across the visual field, from nearby structures to distant trees at the horizon. The warehouse environment was an enclosed space comprising corridors lined by shelves with boxes and other items. In general, surfaces were located at closer distances to the camera and depth varied over a smaller range compared to the neighborhood environment.

At the start of each video, the camera was positioned in the simulated environment at a randomly selected height between 1 and 5 m. The camera then travelled along five connected linear segments each lasting 30 frames. The heading direction for each segment was determined by selecting a pixel within the camera image of the scene at random toward which to move (see Figure 3). Camera speed was fixed within each segment and varied randomly across segments between 1 and 20 m/s. Between segments, the camera rotated to the new heading directions and accelerated or decelerated to the new speed over three frames. As a result, each video lasted 162 frames (or equivalently, 5.4 s at 30 fps). The field of view was 90° H x 90° V. Videos were manually reviewed before inclusion in the dataset to ensure that there were no collisions with objects or the ground.

**Figure 3:**
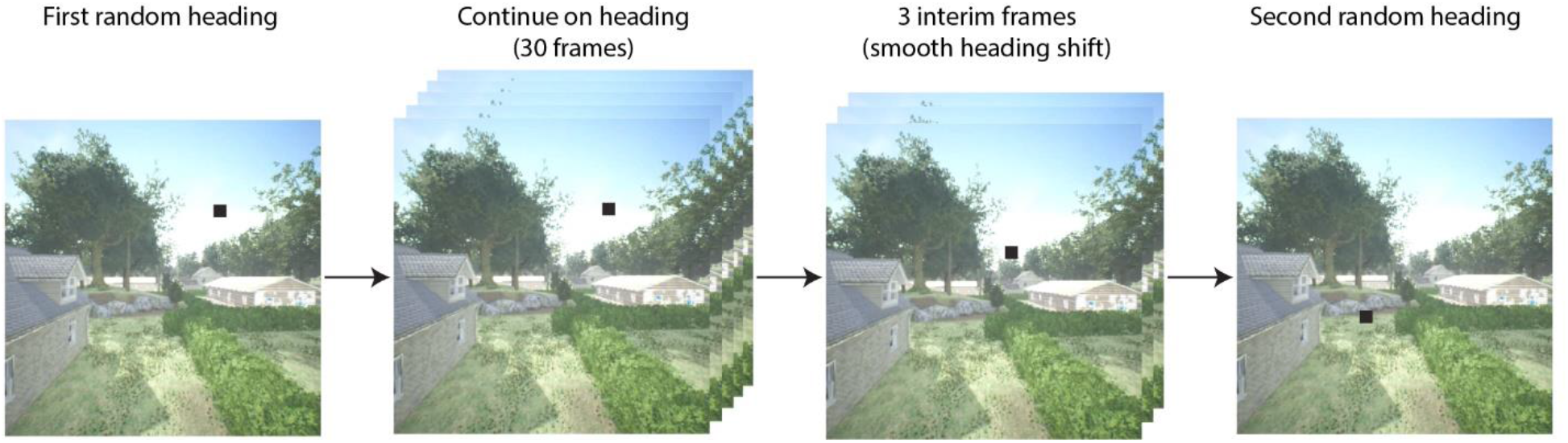
Example of translational segment selection. After an initial heading direction is randomly selected, the camera translates in that direction for 30 frames. Subsequently, a second random heading is selected, and three interim headings are linearly chosen between the first and second heading. After the three-frame transition, the camera translates towards the second heading for 30 frames. This process is then repeated for segments 3, 4, and 5 in each video.

For both experiments, the resolution of the videos was 512×512 pixels and optic flow was estimated using the Farneback method. In the CD model, area MT was set to have seven speed layers, each attuned to a different speed and with a resolution of 256×256 pixels. Model area MSTd had a resolution of 64×64 pixels, meaning the maximally active MSTd neuron specified a heading estimate covering 1.4° of the square, 90° field of view. Downsizing of the spatial resolution balanced model runtime against precision of the heading estimate, which is robust for a range of resolutions but degrades at resolutions much below 64×64 pixels. Model runtime is impacted by the resolution of both model areas but MSTd resolution has a greater effect, as each MSTd unit represents a template cell (e.g. a radial cell) with the pattern centered on that pixel location. Each model template cell necessitates a convolution of the corresponding optic flow template with the feedforward signal from MT. Thus, increasing MSTd resolution rapidly increases the number of calculations necessary. These parameters were selected after preliminary testing revealed that increasing the MT and MSTd spatial resolutions and number of MT speed layers had a negligible effect on model performance.

## Results

### Experiment 1: Comparing dynamic and static tuning

Figure 4a shows the mean absolute heading error per frame averaged across videos for the Dynamic, Static – Uniform, and Static – Aggregate conditions. Absolute heading error was measured by calculating the angle in the 2D image plane between the model estimate and the ground-truth heading direction on each frame (see <u>Movie 4</u>). The four sudden transitions in the heading error time series shown in Figure 4a are a consequence of the abrupt shifts in heading direction between segments. Segments 2 through 5 were preceded by three transition frames, during which heading was rapidly changing from the previous direction to the new one. Heading error surged during the transition period, then gradually dropped due to temporal dynamics within the competitive dynamics model – recurrent competitive interactions among units in MSTd enable smooth, non-instantaneous changes in estimates from the previous to current heading even in the face of abrupt changes in optic flow input (Layton & Fajen, 2016b). The first segment of each video is unique in that there is no existing neural activity from previous stimuli for activity due to the new heading to overwhelm, so the heading estimate quickly stabilizes. For this reason, the first segment was excluded from the following analyses.

**Figure 4:**
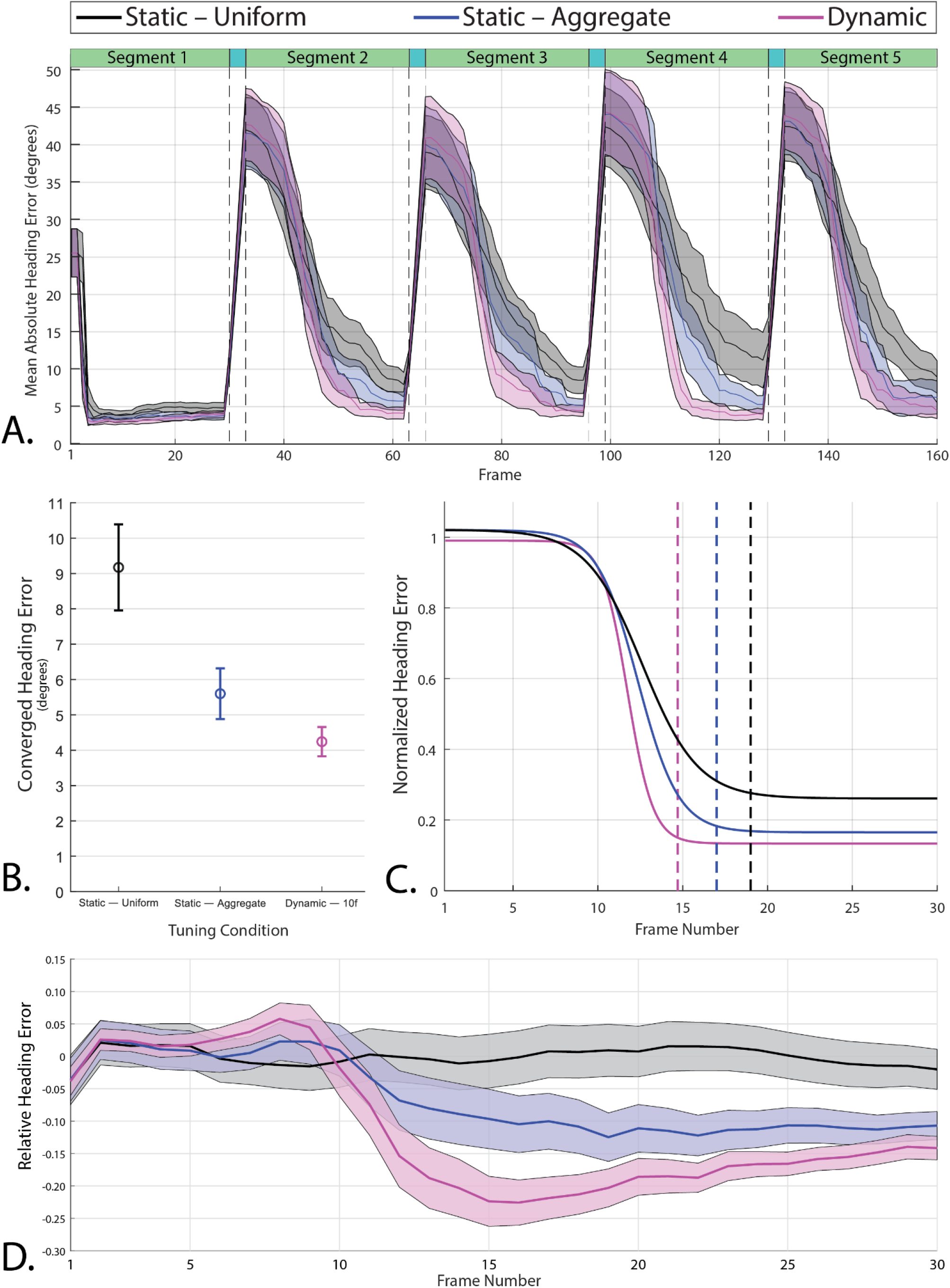
(a) Mean absolute heading error across all 60 videos in the Static-Uniform (black), Static-Aggregate (blue), and Dynamic (magenta) conditions. The shaded regions represent the 95% confidence interval of the mean heading error in each frame. The dashed vertical lines indicate the beginning and end of the 3-frame transition to a new heading prior to the 2nd through 5th segments. (b) Mean converged heading error in segments 2-5 (N=240) with 95% confidence intervals. (c) Best-fitting logistic curves for the heading error in segments 2-5 of each condition (N=240), normalized to the magnitude of the heading shift for that segment; i.e. heading error starts around 1 regardless of whether the initial heading differed from the previous heading by 1 degree or 80 degrees. The vertical dashed lines indicate the frame at which each curve reaches 98% of its range, which provides a measure of each curve’s convergence onto its final heading estimate. Note that the best - fitting curve for the dynamic attunement condition converges more quickly to a lower and therefore more accurate heading error than the static conditions. (d) Mean heading error per frame of segments 2-5 (N = 240) relative to the best-fitting curve of Static-Uniform condition.

We compared the Dynamic, Static – Uniform, and Static – Aggregate tunings on two measures of heading estimation, both of which were extracted from the time series of absolute heading error. The first measure was the converged heading error, which was based on the heading error on the final frame of segments 2-5. Dynamic attunement was more accurate than both static conditions resulting in a mean converged heading error of 4.24° (95% CI [3.83, 4.66]) which was lower compared to both the Static – Uniform (M = 5.60°, 95% CI [4.88, 6.32]) and Static – Aggregate (M = 9.17°, 95% CI [7.96, 10.39]) conditions (Figure 4b). Heading error was also more consistent across segments and trials in the Dynamic condition.

Although the accuracy of model heading estimates relative to human estimates is not directly relevant to the main focus of this study, it is worth noting that with dynamic attunement, model estimates are only slightly less accurate than those of human subjects under similar self-motion conditions. During linear translation through a static environment, human heading judgments are accurate to within 1-2° (Foulkes et al., 2013; William H. Warren et al., 1988), which is ~2-3° more accurate than model estimates. However, in studies of human heading perception, the range of simulated self-motion directions is typically restricted to a two-dimensional plane and subjects are instructed to judge heading direction within that plane (i.e., the azimuth of heading). This contrasts with the conditions used in the present study, in which self-motion direction varied in both azimuth and elevation and the primary measure of heading accuracy was the angular difference between the estimated and actual heading in two dimensions. When heading error is decomposed into its two components, we find that with dynamic tuning, mean converged heading error was 2.44° (95% CI [2.13, 2.76]) along the azimuth and 2.99° (95% CI [2.64, 3.34]) along the elevation axis, which is within 1° of human-level performance. Furthermore, Farneback optic flow estimation is imperfect, introducing noise into the model’s estimate. For human observers, adding noise to the vector directions in the optic flow field resulted in degraded heading accuracy (Warren et al., 1991).

The second measure of performance that we considered was latency, which we define as the number of frames it took the model to converge to a stable heading estimate after each transition. We estimated latency by first normalizing the heading error time series for each individual segment (excluding the 1^st^ segment) of each video such that the heading error on the first frame was 1. This allowed us to parse out the variance due to differences in the magnitude of the heading shift between segments, which varied widely from a few degrees of visual angle up to about 80 degrees. We then fit a logistic curve to each time series, then averaged the curve parameters to determine the average logistic curve for each condition. We defined the heading convergence as the number of frames that was needed for the best-fitting curve to drop by 98% of the difference between the upper and lower bounds of the logistic function. We found that heading error decreased in fewer frames with Dynamic tuning (14.7 frames) compared to both Static – Aggregate (17.0 frames), and Static – Uniform (19.1 frames) (see Figure 4c).

To better visualize the differences between the three tuning conditions, we calculated for each segment the difference between the actual heading error in each tuning condition and the best-fitting curve in the Static – Uniform condition. That is, we subtracted the curve that best fit the heading error time series (segments 2 through 5 of Figure 4a) in the Static – Uniform condition from the actual heading error time series on each trial across conditions. The curves in Figure 4d show the mean and 95% CI of this difference for each frame. This way of representing the data highlights how the improvement in the heading estimate due to dynamic tuning evolves over time while controlling for the size of the heading shift and other sources of variance. Heading error is similar across tuning conditions for the first 8-10 frames but then begins to decrease more rapidly in the Dynamic and Static – Aggregate conditions. By Frame 15, the mean heading estimate in the Dynamic condition is more accurate than the heading estimate in the Static – Uniform condition by about 22.4% of the heading shift magnitude, which is considerably better than the improvement with Static – Aggregate tuning (~9.7% on Frame 15). Over frames, the difference between the three conditions shrinks, suggesting that the primary benefit of Dynamic tuning is that it allows the model to reach an accurate heading estimate in fewer frames.

### Experiment 2: Comparing dynamic time horizons

Experiment 1 demonstrated that with dynamic tuning based on the distribution of optic flow speeds over the past 10 frames, the model converges toward a more accurate heading estimate in less time. In Experiment 2, we examined whether the benefits of dynamic tuning depend on the number of previous frames used to build the optic flow distribution. Specifically, we compared model performance with 10 frames against performance with 1 frame and 30 frames. As shown in Figure 5, the manipulation of time horizon with the range that was tested had negligible effects on both converged heading error and latency. Some small differences were observed in the latency of the heading estimate with a 30-frame time horizon (Figure 5b-c). On some frames in the single frame time horizon condition, the derived speed tuning curves collapsed to identical values as the observed flow was mostly one value, producing erratic activity in this limit case. These outlier issues aside, the model performed similarly across the three time horizon conditions that were tested.

**Figure 5.**
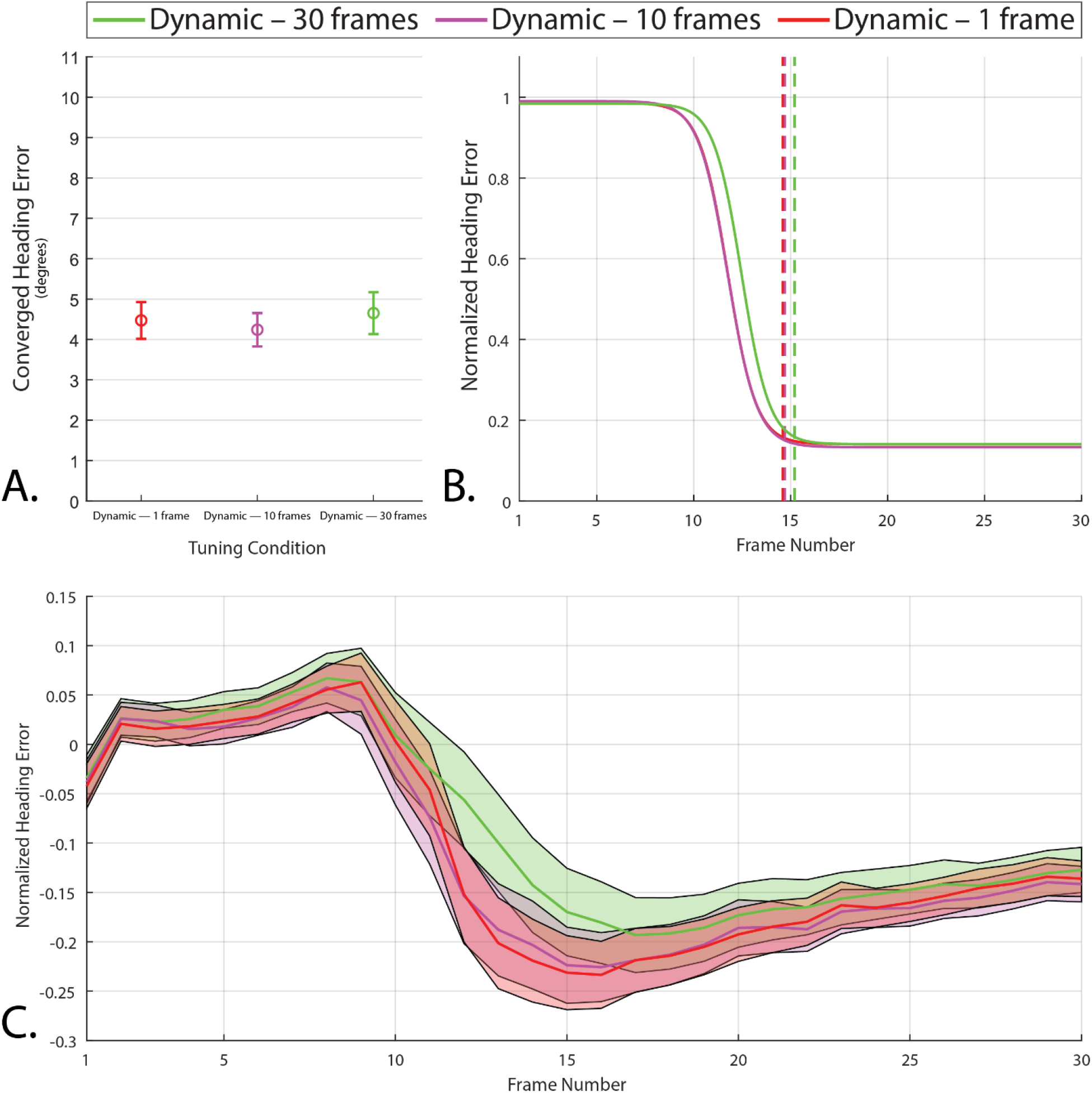
Graphs calculated in the same manner as Figure 4 based on the model with dynamic tuning using time horizons of 1, 10, and 30 frames. Note the absence of major differences among these conditions in the converged heading error (a) or frame of convergence (b), although the analysis of normalized heading error over frames (c) reveals a small difference in the 30-frame time horizon condition on frames 12 through 20.

## DISCUSSION

Incorporating dynamic efficient sensory encoding into the competitive dynamics model improved the accuracy and responsiveness of the heading estimate, bringing the average error closer to human performance on similar tasks. Despite these gains, one might wonder how plausible such a mechanism is in the human visual system, and how downstream areas of the visual system might handle tuning that continually fluctuates based on recently detected stimuli. Here we address both of these issues in turn.

### Biological plausibility

Although many forms of visual adaptation have been observed throughout the early visual system, there remains the question of whether speed cells in MT adapt to recently detected optic flow as well as whether that adaptation occurs on a timescale similar to DESE as implemented here. Let us first consider evidence that the tuning curves of many MT neurons are dynamic rather than static. Dynamic shifts in the directional selectivity of MT neurons have been demonstrated to solve the aperture problem: tuning shifts from the direction of motion perpendicular to a contour in their receptive field to encoding the true direction of motion over a 60 ms period of exposure (Pack & Born, 2001). There is also evidence that surround modulation of MT neurons is stimulus-dependent, with some surrounds becoming either excitatory or inhibitory depending on the stimulus (Huang et al., 2007). Furthermore, the encoding of direction of motion of multiple transparently moving stimuli shifts over time from an averaged response to encoding separate directions (Xiao & Huang, 2015). At the behavioral level, humans have exhibited improved discrimination of recently seen optic flow speeds produced by walking and standing still (Durgin & Gigone, 2007). Taken together, these findings demonstrate that adaptation to recently presented optic flow and other visual stimuli occurs at least to some degree in MT as well as earlier areas.

There is also evidence that adaptation in MT occurs over periods of time that correspond to the dynamic conditions tested in the present study. The highest performing condition from both experiments, the dynamic 10-frame time-horizon condition, simulates adaptation on a timescale that is dependent on the framerate of the input video. Assuming video at 30 frames per second, a 10-frame time horizon equates to 333 ms. By comparison, adaptation has been observed in the activity of macaque V1 orientation neurons after a 400 ms adaptation period (Gutnisky & Dragoi, 2008), and human adaptation to recently detected optic flow was produced after exposures to motion patterns for 1.25-1.5 s (Durgin & Gigone, 2007), which is comparable to the dynamic 30-frame time-horizon condition. These findings lend support to the existence of adaptive mechanisms that operate on timescales similar to those that were simulated in the present study.

Even if DESE does not simulate the exact, low-level mechanisms of adaptation in biological organisms, we contend that it captures the effects of adaptation in a functional sense. Adaptation serves to improve perceptual discrimination via efficient encoding of the stimulus distribution, which is precisely what DESE achieves. In the same way that the rate model of the neuron does not specify or attempt to capture the dynamics of neural activity at the lowest level (Brette, 2015), dynamic application of efficient sensory encoding provides a means of capturing the consequences of adaptive mechanisms on the feedforward signal without simulating the exact mechanism of adaptation. For example, there is some evidence that MT neurons in fact do not adapt their sensitivities to recently detected stimuli, but rather that later brain areas adjust their weighting of various MT neurons as they become more or less relevant to the current stimulus (Liu & Pack, 2017). If this were the case, DESE could be thought of as an approach to simulating in fewer computations the behavior of many statically tuned neurons. From this perspective, the tuning curves calculated under DESE are the subset of neurons from the much larger population that have the most relevant activity (at that point in time) for the property being encoded. These tuning curve selections are optimal with respect to the number of neurons being simulated, assuring the most efficient encoding given computational constraints or, alternatively, capturing the properties of the feedforward signal of a large population of neurons with a small number of simulated ones. Whether the mechanisms behind perceptual adaptation lie in MT or beyond, the resulting effects can be captured using dynamic efficient sensory encoding as an abstraction of adaptation.

### Downstream interpretability

One intriguing implication of dynamic tuning in MT is that the signal sent to the downstream model area (i.e., MSTd) reflects not only changes to the input to MT but also by shifts in the tuning curves. As such, a particular response from a given speed cell may result from motion at one speed at one point in time and a different speed at a later point in time. This raises the question of how the downstream area makes use of that speed cell’s activity to generate an accurate heading estimate when that activity could have resulted from many different inputs.

We hasten to point out that the answer does not require giving the downstream area (MSTd) access to the adapted tuning parameters of MT cells. As argued by Hosoya et al. (2005), there is no need to communicate the current state of adaptation in one area to downstream areas. The brain does not construct a perfectly veridical reproduction of light on the retina but instead detects information that is useful and relevant to behavior. In the present study, the relevant perceptual variable is heading direction and the relevant visual information is carried in the directions rather than the magnitudes of optic flow vectors. For pure translational self-motion, heading is specified by the position of the FoE in the global radial flow pattern, which is invariant over changes in the speed of individual motion vectors. Indeed, human heading estimates during translational movement are unaffected by the addition of noise to flow vector magnitudes but significantly impaired by noise added to vector directions (Warren et al., 1991). This is adaptive because variations in flow vector speed could result from variations in depth or self-motion speed, even if self-motion direction is constant. In contrast, the direction of motion vectors depends on self-motion direction alone. Likewise, in this formulation of the CD model, the template match between the observed optic flow and each radial cell in MSTd is determined largely by the match in vector directions while differences in speed have a minimum impact. Just as the information about heading is largely invariant over flow-vector magnitude, so is the activity of individual radial cells in MSTd. Taken together, although dynamic tuning affects the encoding of optic flow speed in MT, the signals that carry the information relevant to heading estimation are left intact and the impact on MSTd cell activity is minimal.

This explains why dynamic tuning does not degrade heading estimation, but it does not explain why performance improves. The key insight is that dynamic tuning allows the model to be more sensitive to all the optic flow vectors present. By continually shifting tuning curves to detect a larger portion of the current optic flow, more of the available information about heading direction can be detected. As such, even if dynamic tuning-related changes distort the absolute speed signal, the overall improvement in sensitivity to task-relevant information more than compensates for the loss.

It is also worth noting that some information about flow vector speed is preserved in the signal to MSTd. In speed cells governed by DESE, each speed layer captures an equal portion of the probability mass of the stimulus distribution and their ordinal relation remains constant. That is, the speed layer that responds to the slowest speeds is always the slowest speed layer, no matter what distribution is being used for attunement. As such, although the absolute magnitude of optic flow to which a speed cell is sensitive is continually shifting, each speed cell always represents a relatively constant percentage of the stimulus distribution. For example, the slowest speed cell layer might represent the presence of the slowest 10% of speeds in the receptive field of an active neuron. In this sense, DESE converts the signal from speed cells into relative units, where the absolute magnitude of the optic flow is lost but the magnitude relative to recently experienced optic flow is preserved. This could be useful for estimating properties other than heading that may rely on the magnitude of optic flow.

## Conclusion

Dynamic efficient sensory encoding offers a principled approach for capturing adaptation in early visual areas. By adjusting the tuning curves of simulated neurons based on the distribution of recently experienced stimuli, DESE produces the optimal encoding of that distribution by a finite number of neurons which results in improved sensitivity to task-relevant information. Implementing this adaptive mechanism in the speed cells of the competitive dynamics model led to measurable gains in performance, decreasing the error and shortening the latency of heading estimates derived from model area MSTd activity. Although some aspects of the approach are specific to the model and task used in the present study, dynamic efficient sensory encoding is more generic and could also be useful to optimize the encoding of stimulus variables in other domains. Because it relies only on the distribution of recently experienced stimulus values, it does not assume prior knowledge of the statistics of the environment, thereby capturing how biological systems and neural models alike could rapidly adapt to unfamiliar environments.

